# Microtubule-mediated cell shape regulation contributes to efficient cell packing in *Arabidopsis thaliana* cotyledons

**DOI:** 10.1101/2023.05.16.540958

**Authors:** Timon W. Matz, Ryan C. Eng, Arun Sampathkumar, Zoran Nikoloski

**Affiliations:** Bioinformatics, Institute of Biochemistry and Biology, University of Potsdam,14476 Potsdam, Germany; Systems Biology and Mathematical Modelling, Max Planck Institute of Molecular Plant Physiology, 14476 Potsdam, Germany; Plant Cell Biology and Microscopy, Max Planck Institute of Molecular Plant Physiology, 14476 Potsdam, Germany; Max Planck Institute for Plant Breeding Research, 50829 Cologne, Germany

## Abstract

Recent advances have started to uncover the mechanisms involved in organ and cell shape regulation. However, organizational principles of epidermal cells in different tissues remain poorly understood. Here, we show that polygonal representations of cotyledon pavement cells (PCs) in *Arabidopsis thaliana* exhibit increasing irregularity in side lengths and internal vertex angles during early stages of development. While the shape of PCs in cotyledons is more complex than that of cells in the shoot apical meristem (SAM), the polygonal representations of these cells share similar irregularity of side length. Comparison of the surface cell area with the area of the regular polygons, having optimally spaced tri-cellular junctions, reveals suboptimal junction placement for coverage in cotyledons and SAM. We also found that cotyledons show increased packing density compared to the SAM, indicating that PCs forgo coverage of larger areas to potentially increase tissue stability. The identified shape irregularity and cell packing is associated with microtubule cytoskeleton. Our study provides a framework to analyze reasons and consequences of irregularity of polygonal shapes for biological as well as artificial shapes in larger organizational context.

**Summary:** We provide a polygonal cell representation in a tissue context and use it to draw conclusions about cell packing in epidermis of *A. thaliana* cotyledon.

## Introduction

Plant leaves show immense diversity in their shape between and within species (Vőfély et al., 2019). As a result, it is of great interest to determine if and how the shape of leaves relates and is affected by the shapes of individual cells (McLellan and Endler, 1998; Huang et al., 2018; Vőfély et al., 2019; Byrne, 2022). The leaf epidermis contains three types of cells: pavement cells, guard cells, and trichomes, displaying characteristic differences in their cell shapes (Glover, 2000). Guard cells are composed of cell pairs forming pore-like structures called stomata that enables gas exchange (Vatén and Bergmann, 2012). Trichomes are branch-like structures suggested to protect the leaf from biotic and abiotic stress (Wang et al., 2021). Pavement cells (PCs), as the most abundant cell type in the leaf epidermis, often display interdigitated or puzzle-like cell outlines with local regions of the cell protruding into neighboring cells called lobes, and indenting domains called necks (Panteris and Galatis, 2005; Vőfély et al., 2019). Multiple functions for the complexity of PC shape have been suggested (see review of Liu et al., 2021), with one being that the complex shape of the PCs helps to manage internal turgor-driven mechanical stresses by redistributing them from the central region to the boundaries of the undulated cell domains, thus helping in the maintenance of structural integrity as cells enlarge (Sapala et al., 2018).

Mounting evidence points at the role of the cell cytoskeleton in forming the complex morphology of PCs. For instance, the importance of microtubules (MTs) for PC shape complexity has been known for some time (Panteris et al., 1993) with MTs guiding the deposition of the major load-bearing cell wall component cellulose (Paredez et al., 2006). Quantification of cotyledons treated with the MT depolymerizing drug oryzalin (Baskin et al., 1994) further showed no increase in PC shape complexity over time compared to mock treatment in cotyledons, (embryonic leaf-like tissue) (Eng et al., 2021). Additionally, the double knockout of two MT-associated proteins, CLASP and KATANIN, leads to the abolition of lobes and necks, with the single knockout of KATANIN leading to decreased shape complexity (Eng et al., 2021). The recruitment of KATANIN is known for its MT severing capabilities, especially under blue light that stimulates MT re-organization (Lindeboom et al., 2013; Zhang et al., 2013).

To facilitate the study of molecular players that affect cell shape complexity, tissues have been abstracted in several ways. For instance, tracking points along PCs by individual fluorescent particles has led to the discovery of the isotropic (non-directional) mode of lobe growth (Elsner et al., 2018). The representation of cells by individual nodes in a network of juxtaposed cells in a tissue context has identified that cells with more neighbors are more likely to divide (Gibson et al., 2011; Jackson et al., 2019) and facilitated the development of predictive models for information flow and cell division (Jackson et al., 2019; Matz et al., 2022). This representations has led to the observation that the distribution of number of neighbors is robust for PCs across the whole leaf over time (Carter et al., 2017). In comparison to the leaf tissue, with the interdigitated PCs, the shoot apical meristem (SAM) provides a contrast in cell shape complexity as the meristematic cells are almost polygonal and highly regular (Kwiatkowska, 2004). Yet, the meristematic cells in the SAM also show similarly robust neighbor distributions and are suggested to have heterogeneity within the cell walls (Long et al., 2020).

Different approaches have emerged to quantify interdigitation based on the cells overall shape characteristics: aspect ratio, of a cells bounding box bounding (Kuan et al., 2022), solidity, a ratio of a cell’s convex hull with surface area (Wu et al., 2016), or lobyness, setting the perimeter of the cell in contrast with its convex hull (Wu et al., 2016; Sapala et al., 2018). Other methods go into more detail in identifying lobe and neck positions by either investigating the distances of perimeter to the convex hull (Wu et al., 2016) or specifying lobes and necks by local changes in curvature (Möller et al., 2017). Most recently, GraVis, a network-based approach abstracts the cell outline in a network and identifies points along the contour as lobes or necks based on graph-theoretic concepts also allowing to differentiate plant clades (Nowak et al., 2021). Since the leaf epidermis includes guard cells along with PCs, a recent approach has attempted to consider the properties of both cell types (Brown and Jordan, 2023).

With the recent advances in the amount, availability, and aspect of quality in imaging data (Li et al., 2014), but also the improvement of recent protocols for long-term imaging of whole leaves (Seerangan et al., 2020), sharing and storage of ever increasing amount of imaging data could benefit from abstraction and dimensionality reduction for cellular information. In this context and to bridge the gap between simple shape characteristics and very detailed lobe and neck position, we simplify the cell outline into a tri-way junction spanning polygon thus providing a polygonal cell representation. This representation allows us to combine the two defining characteristics of the polygon, its side lengths and internal vertex angles, using the established generalized irregularity measure, the Gini coefficient (Sen, 1978; Bendel et al., 1989). To address the question of underlying complexity from the spacing of tri-way junctions in PCs, we draw upon existing time-course data of cotyledons and utilize the central region of the SAM as a counterpart to cotyledons due to its simple polygonal shaped surface cells. Using this framework, our findings show that: (1) PC polygonal representations length and angle irregularity increases as cells enlarge and is dependent on MT organization, (2) angle irregularity is higher in PCs compared to cells from the SAM, and (3) epidermal cells in cotyledons and SAM could cover larger surfaces with the same junction spacing, indicating compact packing.

## Results

### Length and angle irregularities are properties that emerge from the proposed polygonal representations

To improve the characterization of resulting polygonal shapes, we determined the polygons irregularity of side length and internal angle, utilizing the established method of the Gini coefficient, with higher values denoting larger irregularity in polygonal shapes (see Methods). To illustrate the two Gini coefficients, we ordered artificial shapes (e.g., hexagon, star, and gecko polygons) based on the two irregularity measures (Fig. 1A). We observed different orderings based on the selected property, i.e. polygonal side length or internal angle, with the hexagonal shape being perfectly regular for length and angle, and the star showing regular lengths, but irregular angles. Visualization of the Gini coefficient of length versus angle over different shapes further highlights the importance of considering both properties since they capture different information about polygonal shapes (Fig. 1B).

**Figure 1.**
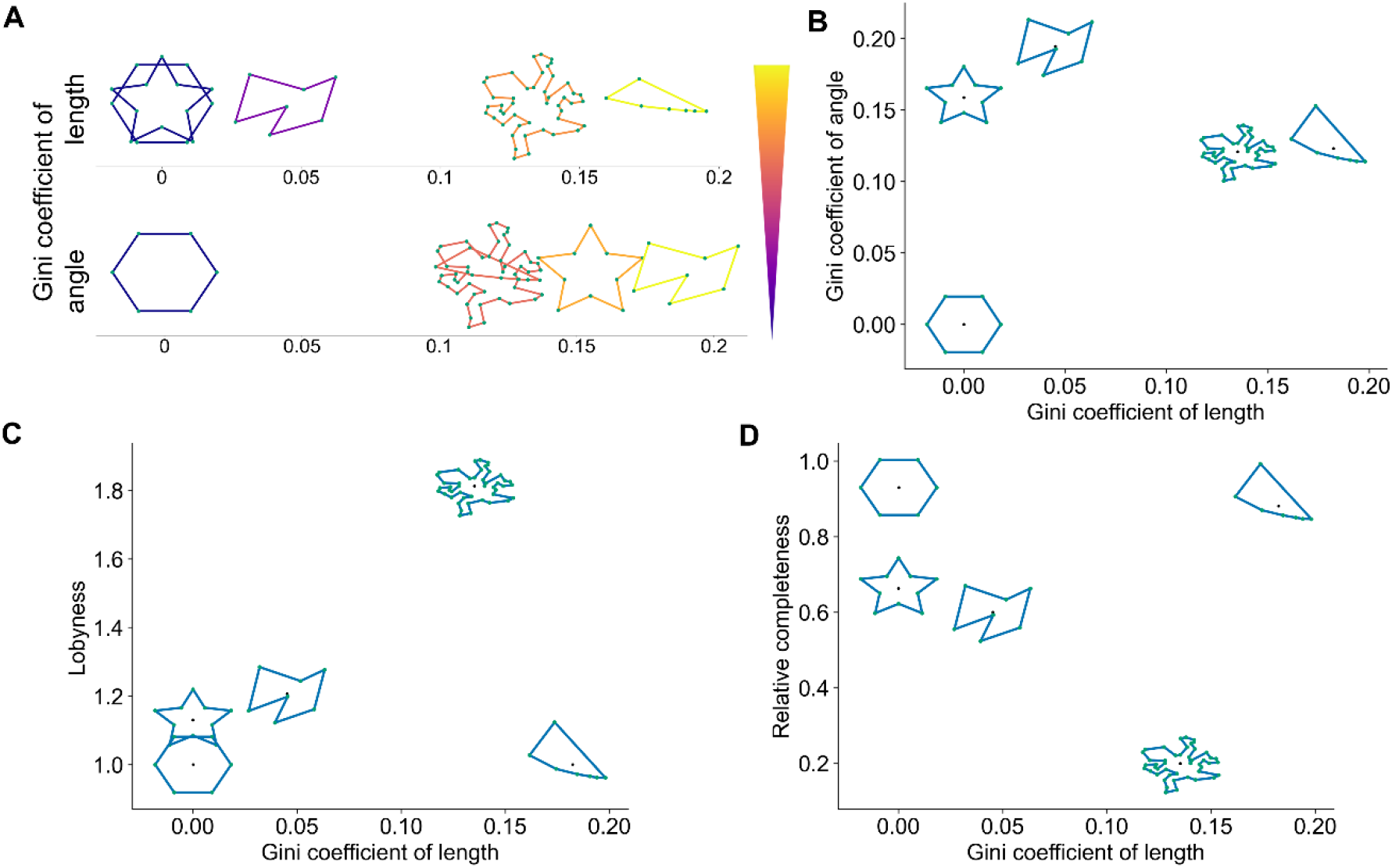
Comparison of length and angle irregularity, lobyness, and relative completeness for artificial polygonal shapes. (A) Visualization of artificial shapes, with color-coded edges and vertices in green, ordered based on Gini coefficient of length (top) and angle (bottom). (B) Association between the Gini coefficients of length and angle (black dot) for different artificial shapes (blue). Association of Gini coefficient of length with (C) lobyness and (D) relative completeness for the artificial shapes shown in panel A.

To put the irregularity into context with other shape characteristics, we compared the Gini coefficient of length with lobyness (Sapala et al., 2018) and relative completeness (Nowak et al., 2021) for the artificial shapes (shown in Fig. 1A). Here, we observed no association between length irregularity and lobyness or relative completeness (Fig. 1C,D). This indicates the different capabilities of length and angle irregularity to analyze internal polygonal properties, and lobyness and relative completeness to identify overall shape characteristics.

### Increased angle irregularity in cotyledons

To apply the irregularity measures on real life examples, we choose the complex shaped pavement cells of five-day old cotyledons and compared them against the surface cells of the SAM from the central region (Matz et al., 2022), which are almost polygonal in nature. We abstracted the cell outline into polygonal representation by creating a tri-way junction spanning polygon, with vertices corresponding to tri-way junctions to calculate the Gini coefficient from the polygonal representations side length and internal vertex angle (Fig. 2D). We displayed the outline and polygonal representation of cells from the whole cotyledon and the central region of the SAM, selecting five cells and coloring their polygonal representation based on length and angle irregularity (Fig. 2A,B). We quantified the irregularities of both tissues, finding similar length irregularities for the polygonal representations despite the more complex outlines of PCs; furthermore, the angle irregularity is increased in cotyledons compared to the SAM (Tukey HSD Test with *post hoc* ANOVA, p-values < 0.05, Fig. 2C). These findings indicate the robustness of relative tri-way junction distancing across different tissues, while highlighting the difference in relative spatial positioning of junctions.

**Figure 2.**
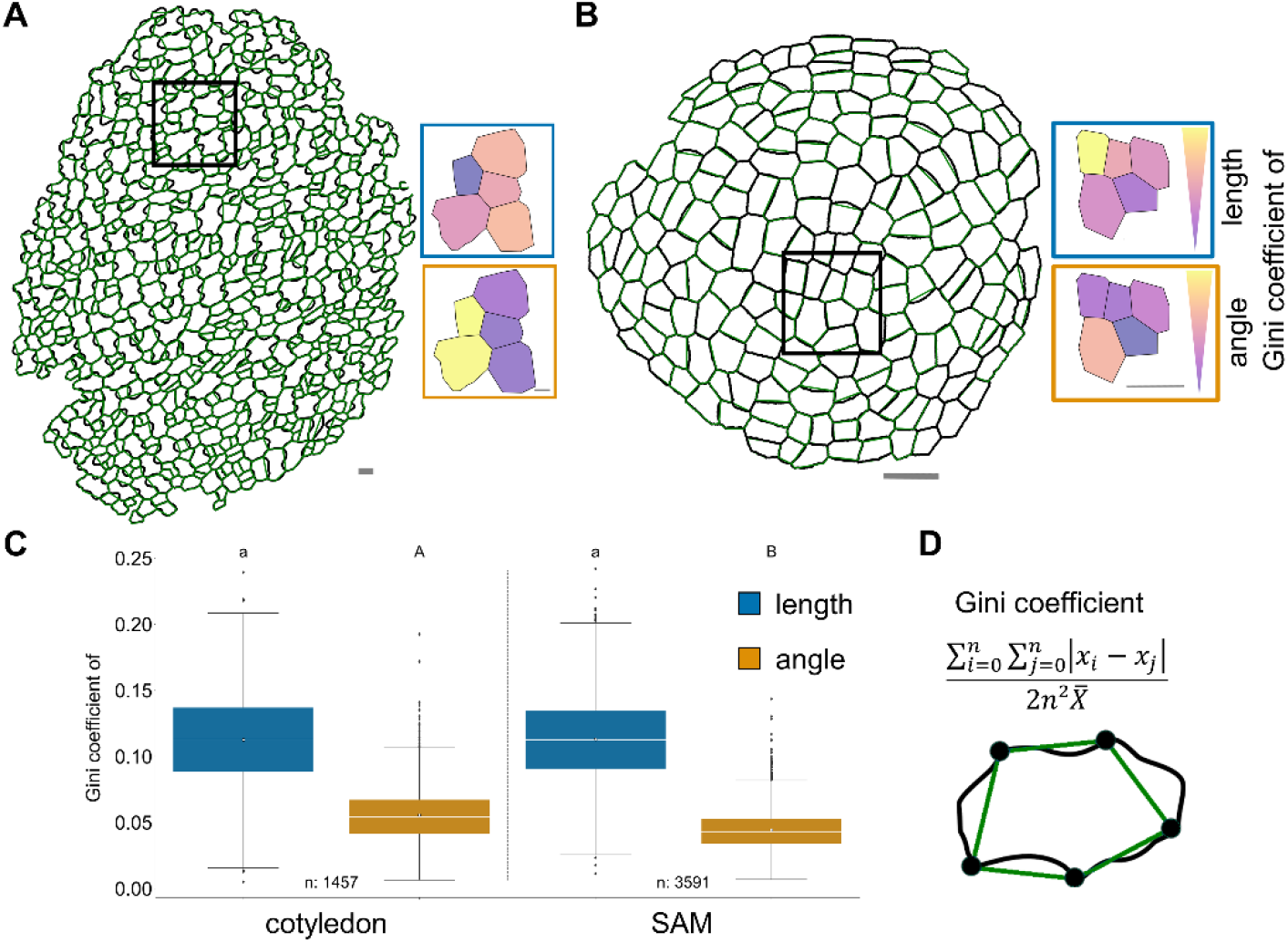
Comparison of polygonal cell irregularities in wild type cotyledon and shoot apical meristem (SAM). Visualization of outline (black) and polygonal cell representation (green) of (A) whole cotyledon (120 hours post dissection) and (B) central region of the SAM tissue. Polygonal cell representations selected from a tissue (black frame) colored based on Gini coefficient of length (blue frame) and angle (orange frame). Scale bars, 10 μm. (C) Quantitative comparison of cotyledon and SAM irregularities (length in blue and angle in orange) with mean (black dot) and median (white line). (D) Illustration of the polygonal cell representation with outline and tri-way junctions (black dots). (E) Mathematical expression for the Gini coefficient, with *x* indicating the length or angle at side or corner *i* or *j*, respectively, *n* denoting the number of sides/corners, and 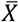 denoting the mean over all *x* values. Different letters indicate significance between groups using one-way ANOVA with Tukey’s pairwise comparison (p-value < 0.05). Number of cells (n) are indicated in the figures.

### Irregularity of polygons is time-dependent and associated with microtubules (MTs)

Next to determine the temporal behavior of polygonal irregularities in cotyledons, we applied the Gini coefficients on the polygonal representations of developing wild type (WT) PCs tracked for a period of four days (Eng et al., 2021) (Fig. 3A,B). As noted above, this visualization clearly demonstrates the independence of the two properties used (Fig. 3B). We observe that during early stages of PC development, polygon irregularity as measured by Gini coefficients of length and angle increased as cells enlarge (Fig. 3C). This indicates heterogeneity in directional expansion of different segments of the cell based on the position of the tri-cellular junctions in developing PCs. Given that MTs play a crucial role in regulating pavement cell shape complexity, we further evaluated developing PCs of oryzalin treated cotyledons in which MTs are completely depolymerized, as well as the *ktn1-2*, a mutant known to have defects in MT organization (Eng et al., 2021). Our analysis indicated that complete abolishment of MTs using oryzalin resulted in no significant changes to the polygon irregularity (Tukey HSD Test with *post hoc* ANOVA, p-values ≥ 0.05, Fig. 3C,D). In 96 h old *ktn1-2* cotyledons, we observed an increased length irregularity and a significantly decreased angle irregularity compared to the WT at 96 h (Tukey HSD Test with *post hoc* ANOVA, p-values < 0.05). Our findings highlight the importance of proper MT organization in contributing to the development of underlying cell complexity in the junction positioning based on the irregularities of the polygonal cell representation.

**Figure 3.**
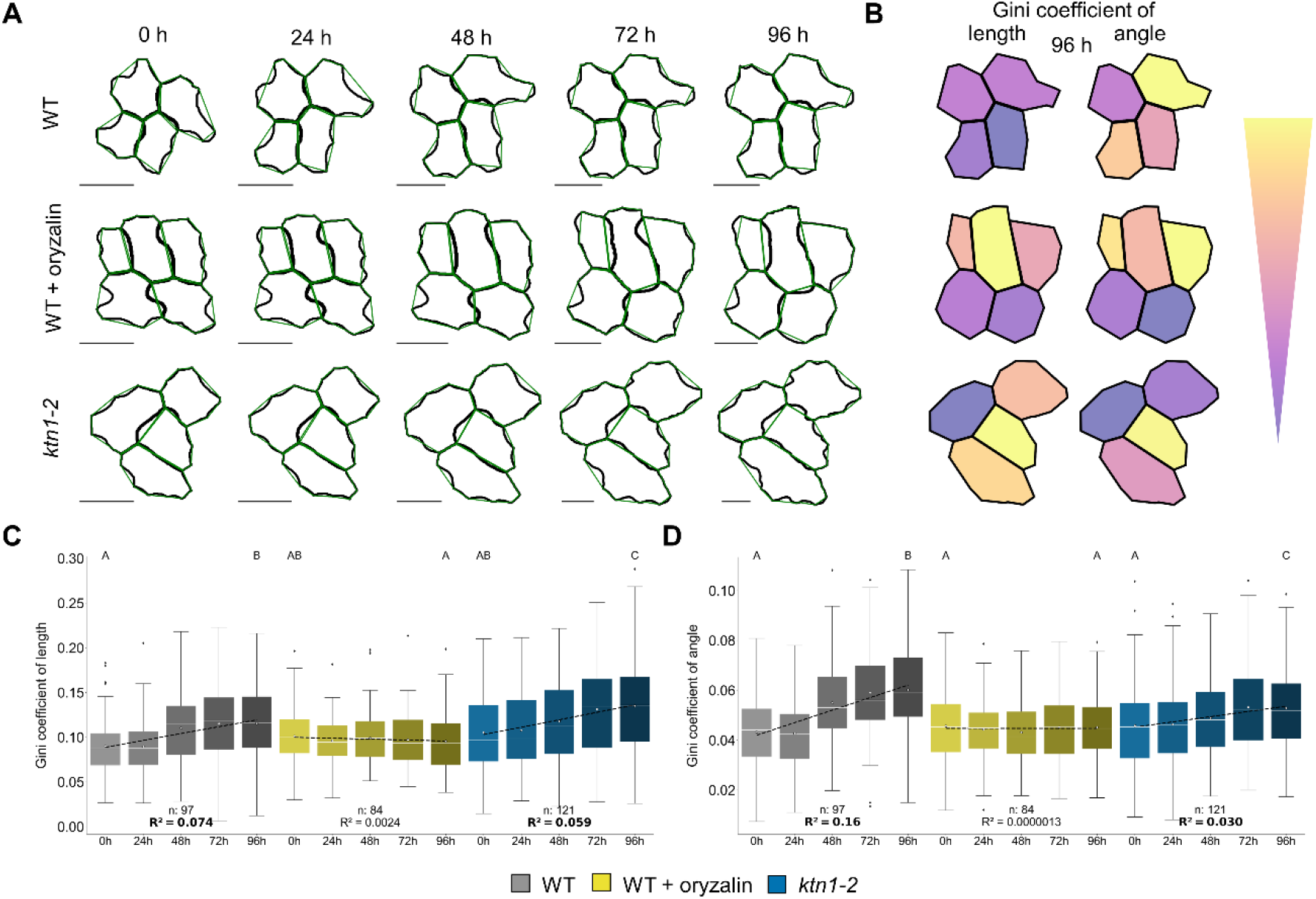
Time-series of polygonal cell length and angle irregularity in cotyledons of different genotype combinations. (A) Outlines (black) and polygonal cells (green) of wild type (WT), WT treated with oryzalin (WT + oryzalin), and *ktn1-2* tracked over 96 hours post dissection (h) in 24 h time steps. Scale bars, 20 μm. (B) Polygonal cell representations are colored based on the Gini coefficient of length and angle for the epidermal tissues of different scenarios at 96 h. Quantification of Gini coefficient of (C) length and (D) angle of different scenarios (WT: gray, WT + oryzalin: yellow, *ktn1-2*: blue) over time (later time points being darker, mean circle, median white line), along with linear regression (dashed) line and R^2^ (significant values with p-value < 0.05 in bold). Non-overlapping letters between groups indicate significance from one-way ANOVA with Tukey’s (p-value < 0.05). Number of cells (n) are indicated in the plots.

### Epidermal cell shape allows for compact packing of cells

To further build on the idea of the polygonal cell shape, we calculated the ratio of the polygonal and regular polygonal area representation with the original cell area to identify the goodness of fit between area of polygonal representation and original cell, and determine cells packing density, respectively (Fig. 4A). The regular polygon is defined as the polygon with equal edge lengths having the same perimeter and number of vertices as the polygonal cell representation. Thus, it represents the maximal area of a polygon with the equal junction spacing, and allows us to investigate packing density. We quantified the ratios and found that the polygonal area ratio is close to one for both tissues (1.9 % and 1.2 % lower than one, respectively, while still being significantly different from one using one-sample t-test, p-value < 0.05, Fig. 4B-D). This observation indicates that the polygonal area is a good proxy for the original cell area. We observed the regular polygonal area ratio to be above one for both tissues (one-sample t-test, adjusted with Benjamini-Hochberg) with the cotyledon being significantly higher than the SAM (Tukey HSD Test with *post hoc* ANOVA, p-values < 0.05). These results indicate that complex changes in PC shapes allows for more dense packing of cells in comparison to the meristematic cells of the SAM.

**Figure 4.**
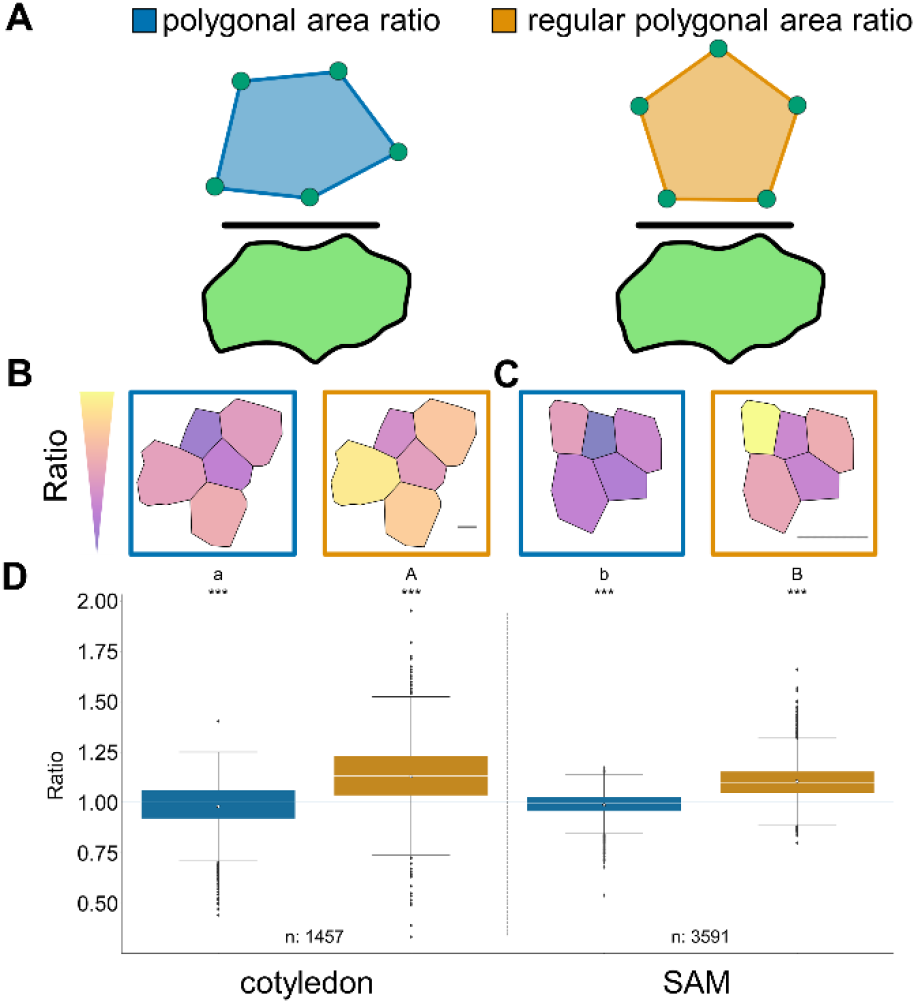
Comparison of polygonal area ratios in wild type cotyledon and shoot apical meristem (SAM). (A) Visualization of polygonal and regular polygonal cell area ratio calculation indicating division of polygonal (blue) and regular polygonal area (orange) by original cell area (green), both polygons having the same perimeter. Coloring polygonal cell representation (of Fig. 2) based of polygonal (blue frame) and regular polygonal cell area ratio (orange frame) from (B) whole cotyledon (120 hours post dissection) and (C) central SAM. Scale bars, 10 μm. (D) Quantification of polygonal (blue) and regular polygonal (orange) area ratios (mean circle, median white line). Ratio of one indicates equal areas of the (regular) polygonal representation and cells. Different letters indicate significance between groups using one-way ANOVA with Tukey’s pairwise comparison with small and capital letters for ratio of polygon to cell area and regular polygon to cell area, respectively (p-value < 0.05). One-sample Benjamini-Hochberg adjusted t-test against one is applied for all groups with p-value: ^***^ < 0.001. Number of cells (n) are indicated in the figure.

### MT rearrangement affects the increase in packing over time

As a next step, we tested the role of MTs in contributing to the packing behavior of cells. We compared the area of the polygonal and regularized polygonal representation with the original cell area (polygonal and regular polygonal area ratios, see above) in WT, WT + oryzalin, and *ktn1-2* cotyledons over time. We observed polygonal area ratios close to one and above one for the regular polygonal area ratio for the selected cells of all scenarios at 96 h (Fig. 5A). We could further support these observations quantitatively finding polygonal area ratios indeed to be close to one for 18 of 20 scenario and time point combinations (one-sample t-test, adjusted with Benjamini-Hochberg, p-value ≥ 0.05, Fig. 5B) with no differences between scenarios and different developmental time points (Tukey HSD Test with *post hoc* ANOVA, p-value ≥ 0.05). Our results indicate that the covered area of the polygonal representation and outline are similar and that the polygonal area can serve as an approximation independent of MTs and the developmental stage.

**Figure 5.**
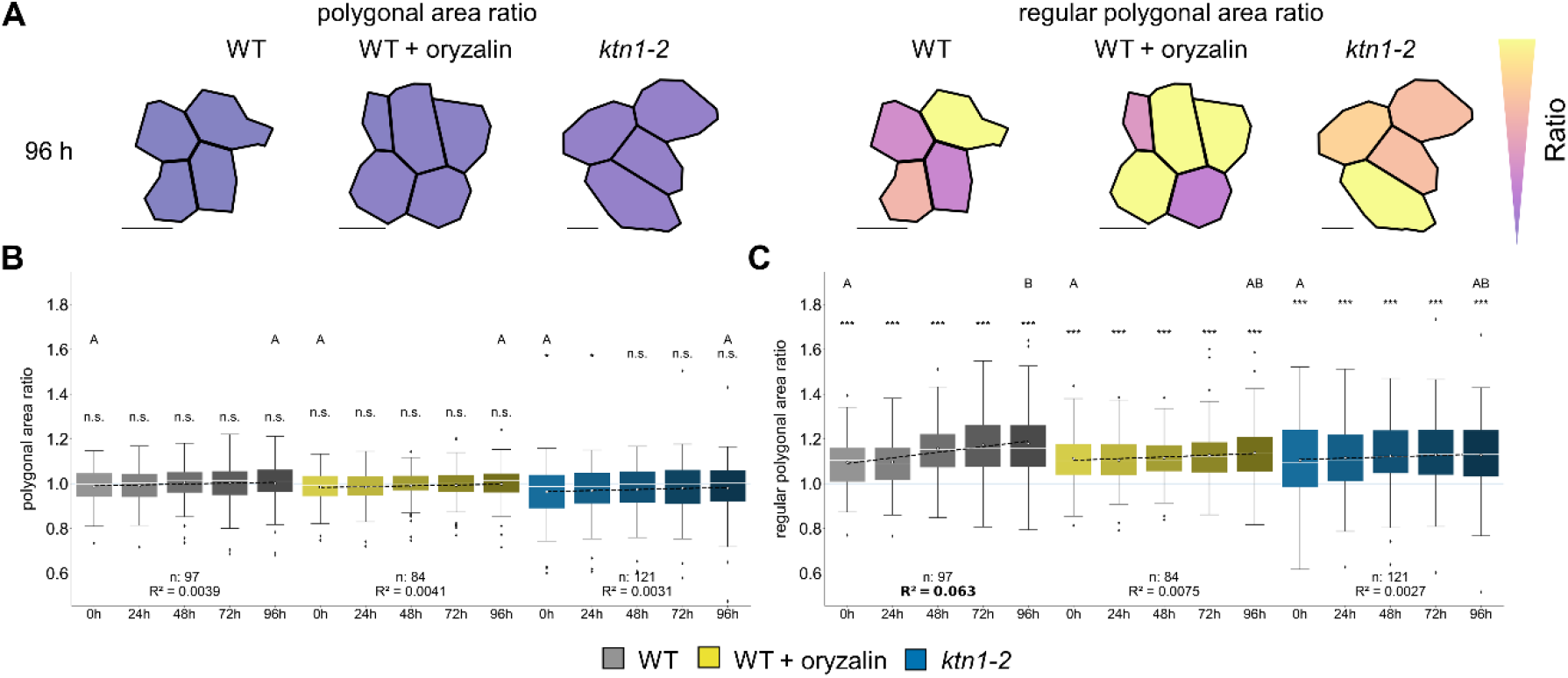
Time-series of polygonal and regular polygonal area ratio in cotyledons of different genotype combinations. (A) Coloring polygonal representation based on polygonal (left) and regular polygonal (right) area ratio for the cotyledon of different scenarios at 96 h. Scale bars, 20 μm. Quantification of (B) polygonal and (C) regular polygonal area ratio (WT: gray, WT + oryzalin: yellow, *ktn1-2*: blue) over time, with later time points depicted as darker, mean denoted by white circle, median by white line; linear regression for the mean ratios in terms of time is depicted by a dashed line, with significant R^2^ values shown in bold (p-value < 0.05). Non-overlapping letters between groups indicate significance from one-way ANOVA with Tukey’s (p-value < 0.05). Number of cells (n) are indicated in the plots.

To further investigate influence of MTs and time on the packing density, we used the regular polygonal area ratio on the same combinations observing all combinations to be above one (one-sample t-test, adjusted with Benjamini-Hochberg, p-value < 0.05). Over four days, we found WT tends to increase the compactness of cell packing, as quantified by the regular polygonal area ratio from significant Pearson correlation and increases of 6.6% from 0 to 96 h (Tukey HSD Test with *post hoc* ANOVA, p-value < 0.05, Fig. 5C). In contrast, we did not find increases in oryzalin treated and *ktn1-2* PCs over time (Tukey HSD Test with *post hoc* ANOVA, p-value ≥ 0.05). With these results, we were able to indicate that MTs contribute to the temporal increase in cell packing density.

## Discussion

Leaves need to resist a variety of stresses, resulting in the need to balance survivability and stability with energy production. The diversity of complex shaped PCs (Vőfély et al., 2019) highlights the importance of interdigitation, with existing overall cell shape and detailed complexity measures aiding in characterization of cell shapes. However, the default measure of cell shape complexity remains lobyness or the number of neighbors, which has been extensively studied in leaves (Carter et al., 2017; Fox et al., 2018; Le Gloanec et al., 2022), and the latter is used as a major property to access cell division models goodness of fit (Farhadifar et al., 2007; Long et al., 2020). Based on the combination of the Gini coefficient, an established measure of irregularity and the concept of the tri-way junction spanning polygonal cell representation, we proposed a framework allowing for finer characterization of irregularities in the tissue context.

Our result of cell shape analysis based on polygon representation reveals that proper organization of MTs influence the displacement of tri-cellular vertices becoming more irregular and adding to the complexity of the cell outlines. Our analysis also reveals that the packing density in form of the ratio of regular polygonal to original cell area is increasing in cotyledons over time. As PCs cover the largest part of the leaves surface and allow for space for stomata to conduct gas exchange, they show more dense packing than compared to the central region of the SAM, hinting at a trade-off in the PCs as they are incentivized to increase the covered surface area (Gonzalez et al., 2012) per cell, which would be possible by more optimally spacing tri-way junctions. With the generalizability of dividing any surface into polygonal representation the irregularity analysis utilizing the Gini coefficient of length, angle, or alternate edge and vertex related properties allows for new ways to interpret regions of interest in a range of different fields.

## Material and Methods

### Image data

We used existing wild type (WT), oryzalin-treated WT (WT + oryzalin), and *katanin1-2* (*ktn1-2*) cotyledon data sets measured over four days data from Eng et al., 2021). Four replicates of the entire adaxial surface of five-day old WT cotyledons were imaged with spinning-disk confocal microscopy using a 60x water objective lens (NA = 1.2). Over-lapping regions of the cotyledon were acquired and stitched using ImageJ (Schindelin et al., 2012) followed by analysis using MorphoGraphX (MGX) (Barbier de Reuille et al., 2015). As a reference tissue, we used shoot apical meristem surface (Matz et al., 2022).

### Data extraction

To extract outlines and tri-way junctions of polygonal representations, we loaded the surface meshes into MGX and extracted the surface area of the cell centers (‘Process/Mesh/Heat Map/ Heat Map Classic’), converted surface into a cell mesh (‘Process/Mesh/Cell Mesh/Convert to a cell mesh’ with the ‘Max Wall parameter’ set to -1), and saved the contour data, and cellular graph representation as ply.-format (‘Process/Mesh/Export/Cell Graph Ply File Save’). We extracted the positions of tri-way junctions from the contour data of the cell mesh (converted from the surface with the ‘Max Wall parameter’ of 0 instead of -1) and the respective surface areas of the polygonal representation as described above. We excluded the stomata and removed small cells below the 25% percentile area threshold of the five-day old cotyledons, to enrich the number of PCs and reduce the number of stomata precursor cells.

### Cellular properties

For the computational analysis, we used python 3.8.1 and the entire code to reproduce the findings is available at https://github.com/matz2532/irregularity_and_polygonal_representation. We automatically extracted the tri-way junction position in each tissue and determined the tri-way junctions of each cell. We verified and added missing tri-way junctions manually (for the data not present in MGX-format). By ordering the tri-way junctions in a clock-wise manner, we extracted the polygonal representation of the selected cells.

We extracted the length of each side (i, j) and the angle between two sides of the polygonal representations, to calculate the irregularity of the tri-way junction spanning polygon using the Gini Coefficient 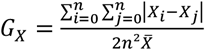 (a value for equality with a range between 0 and 1) (Sen, 1978; Bendel et al., 1989) of the side length and angle for each cell (with *X* either being the side length or internal angle, and 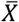 being the mean over all *X*). A *G*_*X*_ of 0 represents a polygon with equal side lengths or angles, and value close to 1 describes a completely irregular polygon. As general shape characteristics, we calculated lobyness (Sapala et al., 2018) and relative completeness (Nowak et al., 2021).

To compare the cell’s surface and polygonal cell area, we computed the ratio of polygonal and regular polygonal cell area (*A*_*regular*_) to surface area. The regular polygon has the same number of sides (neighbors) and perimeter, but all side have the same length 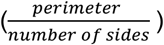 with an area of 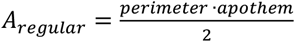.

## Code availability

The entire code to reproduce the findings is available at https://github.com/matz2532/irregularity_and_polygonal_representation

## Acknowledgments

Z.N., T.M., R.E., A.S. acknowledge the support by the project SHAPENET, 031L0177A (to Z.N.) and 031L0177B (to A.S.), of the German Federal Ministry of Education and Research.

## Author Contributions

Z.N. and A.S. designed the research. R.E. plant related experiments. T.W.M. implemented the computational approaches and performed the computational experiments. All co-authors contributed to the final version of the manuscript.

